# NoRMCorre: An online algorithm for piecewise rigid motion correction of calcium imaging data

**DOI:** 10.1101/108514

**Authors:** Eftychios A. Pnevmatikakis, Andrea Giovannucci

**Author notes:** Corresponding author Email address (Eftychios A. Pnevmatikakis).

## Abstract

**Background:** Motion correction is a challenging pre-processing problem that arises early in the analysis pipeline of calcium imaging data sequences. The motion artifacts in two-photon microscopy recordings can be non-rigid, arising from the finite time of raster scanning and non-uniform deformations of the brain medium.

**New method:** We introduce an algorithm for fast Non-Rigid Motion Correction (NoRMCorre) based on template matching. NoRMCorre operates by splitting the field of view into overlapping spatial patches that are registered at a sub-pixel resolution for rigid translation against a continuously updated template. The estimated alignments are subsequently up-sampled to create a smooth motion field for each frame that can efficiently approximate non-rigid motion in a piecewise-rigid manner.

**Existing methods:** Existing approaches either do not scale well in terms of computational performance or are targeted to motion artifacts arising from low speed scanning, whereas modern datasets with large field of view are more prone to non-rigid brain deformation issues.

**Results:** NoRMCorre can be run in an online mode resulting in comparable to or even faster than real time motion registration on streaming data. We evaluate the performance of the proposed method with simple yet intuitive metrics and compare against other non-rigid registration methods on two-photon calcium imaging datasets. Open source Matlab and Python code is also made available.

**Conclusions:** The proposed method and code provide valuable support to the community for solving large scale image registration problems in calcium imaging, especially when non-rigid deformations are present in the acquired data.

## 1. Introduction

Calcium imaging methods enable the monitoring of large neural populations over long periods of time with single neuron resolution. Before addressing specific scientific questions, the analyst needs to pre-process the data and extract the neural signals of interest from the fluorescent microscopy time series images/volumes. The typical calcium imaging pre-processing pipeline consists first of motion correction/image registration of the time series, followed by source extraction, where the different neurons and processes along with their neural activity time series are extracted. In this paper we focus on the motion correction pre-processing step: we introduce an algorithm for Non-Rigid Motion Correction (NoRMCorre), that is suitable for the registration of large scale planar or volumetric imaging data, and we evaluate its performance against state-of-the-art algorithms.

The general field of image registration has a long history and is still very active with many different methods available. In the context of fluorescent microscopy time series data, an algorithm needs to be i) fast since each experiment typically consists of tens of thousands of frames, ii) robust to noise arising from measurement noise and neural variability/activity, and iii) able to deal with non-rigid deformations that occur from natural brain movement and/or slow raster scanning. In several cases rigid translation accounts for most of the motion and fast methods based on template alignment are often used [16, 7, 3]. For dealing with non-rigid motion in the context of calcium imaging data, available approaches include the work of Greenberg & Kerr [6] which is based on the Lucas-Kanade method [9], Hidden Markov Models (HMM) [2, 8] approaches, and block rigid registration [11].

NoRMCorre is based on template alignment and operates by estimating a smooth non-uniform motion field that is applied into different parts of each frame. Our goal is not to take a completely new approach to motion correction, but rather to present and make available a robust alignment method that also combines two important features:

- **Online processing:** The algorithm operates by matching patches of each given frame against a template that is continuously updated based on previously registered frames. As such, it requires access only to the current frame to be registered and the running template, plus possibly a small buffer to store past templates. Consequently it is suitable for online registration of high volume streaming data, a useful feature that can facilitate fully closed loop optical interrogation experiments [12] or compensate for limited amounts of available memory.
- **Fast, non-rigid registration:** The brain is a non-rigid, non-uniformly deformable medium. In modern experimental conditions, with animal preparations locomoting or otherwise moving under fixed or head-mountable microscopes, the brain is subject to elastic deformations. This phenomenon is even more evident as equipment allows for the monitoring of increasingly larger brain areas. Therefore, even when imaging at high speed correction of motion by rigid alignment can be inadequate. NoRMCorre splits the field of view (FOV) into overlapping patches that are registered separately and then merged by smooth interpolation. As such it overcomes the shortcomings of rigid motion alignment without a significant computational cost, thus remaining applicable to large scale datasets. Compared to the other available non-rigid registration methods that split the FOV only along one axis to capture the non-rigid motion caused by the finite speed of raster scanning, NoRMCorre treats all axes uniformly aiming to account for natural brain movement as well.

We present an application to resonant scanning two-photon microscopy data and compare it against other non-rigid image registration methods in terms of speed and performance. To quantify performance we propose three custom metrics. Our results indicate that NoRMCorre achieves state of the art results while operating at a speed not significantly slower compared to template based rigid alignment.

## 2. Materials and Methods

### 2.1. Algorithm Description

#### 2.1.1. Registering a frame against a given template

NoRMCorre can operate in a rigid or piecewise-rigid (pw-rigid) fashion. For rigid registration, every frame is aligned against a calculated template at a sub-pixel resolution using the method proposed by Guizar-Sicairos et al. [7]; the displacement vector is computed by locating the maximum of the cross-correlation between the frame and the template. The cross-correlation is efficiently obtained via fast Fourier transform (FFT) methods, and subpixel registration is achieved at a very moderate computational and memory cost by upsampling the discrete Fourier transform only around the location of the maximum, and then refining the translation estimate.

In the piecewise rigid approach, for any given frame we split the FOV into a set of overlapping patches (*Fig. 1a*) according to user determined dimensions and amount of overlap. Each patch is registered against the corresponding part of the template at a subpixel resolution. Next, each patch is further split into smaller overlapping subpatches with user-defined dimensions and amount of overlap. Similarly, the computed displacement vectors for the set of the initial patches are upsampled to create a smooth motion field. This associates to each of the subpatches a new translation vector that is subsequently rigidly applied to it (*Fig. 1b*). The registered sub-patches are then overlaid to each other and in regions of overlap a weighted average is taken between all the participating patches. The registered frame is also used to update the template in the online scenario as discussed in the following section. A block diagram of the registration pipeline is depicted in Fig. 1c.

**Figure 1:**
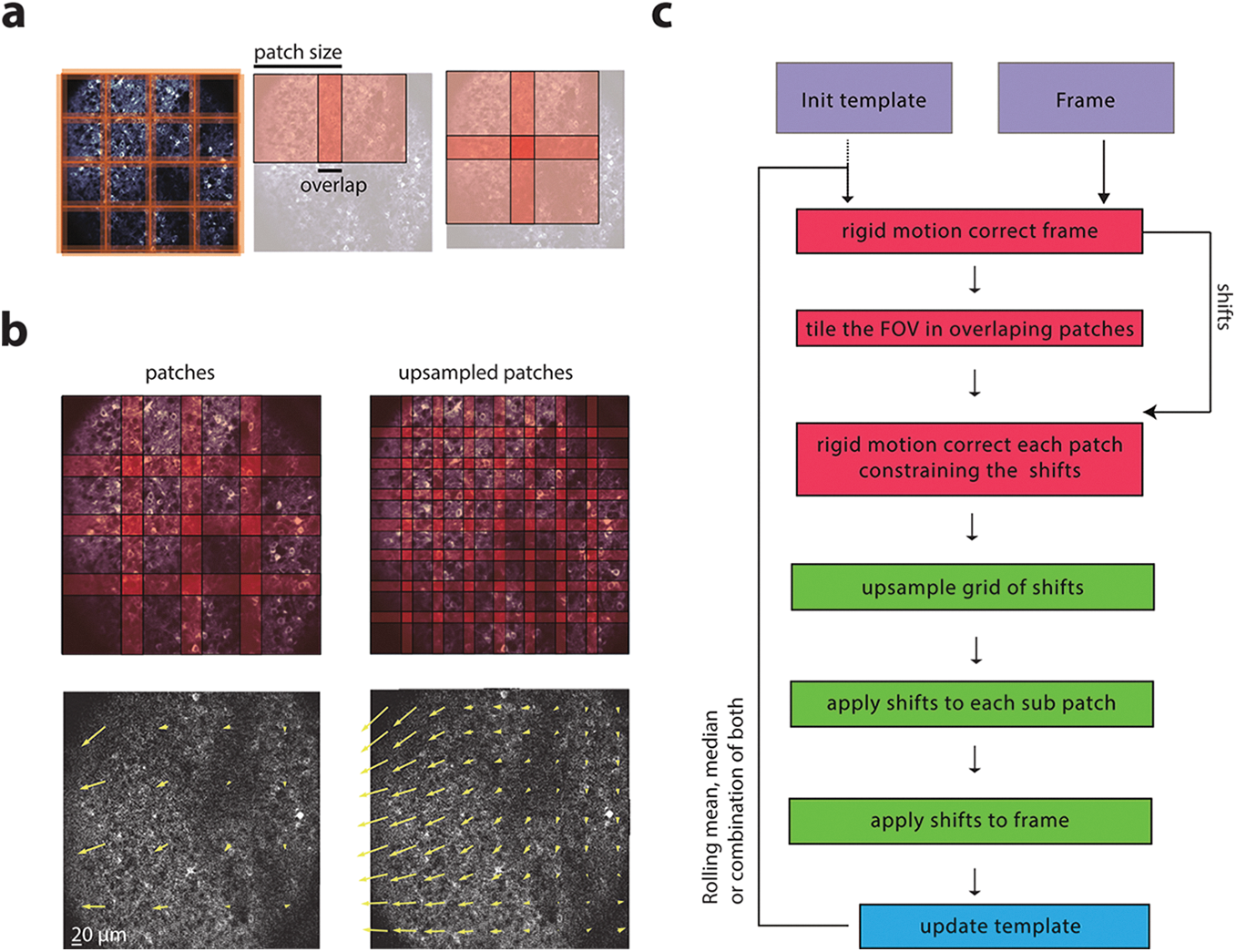
Schematic representation of the proposed algorithm. *a*: Illustration of the scheme used to overlap patches. *b*: *Top.* Illustration of the process of upsampling the shifts. *Bottom.* Visual representation of the motion estimated field on the original (left) and upsampled (right) patches. The yellow arrow’s length represent the direction and magnitude of the motion field. *c*: Pipeline for piece-wise rigid registration of a frame against a given template, and template updating.

#### 2.1.2. Updating the template

The template is updated every *b*_*w*_ frames. Once *b*_*w*_ frames get registered against a fixed template, their average (e.g., mean/median) is computed. These averages are stored in a buffer that keeps at most the last *b*_*p*_ averages. The new template is generated by averaging (e.g., by taking the mean/median) the buffer content. Based on empirical observation, as a default choice, we use the median-of-means to update the template at the end of each minibatch. The template can initialized by computing the median of the first frames (or just the median of a random subset of frames).

#### 2.1.3. Online vs Offline

NoRMCorre is in principle an online and one-pass algorithm since each frame is registered based on the current estimate of the template. However several optimization expedients can be used to improve its performance when data and memory are available. For example to avoid the influence of slow motion trends, especially at the beginning of the motion correction process, we can randomly permute the frames order prior to any registration, or start from the middle time point of the dataset and continue outwards towards the beginning/end. Moreover, when operating in offline mode, the frames within each minibatch that is registered with a fixed template can be processed in parallel, leading to potentially significant computational gains, depending on the available infrastructure.

#### 2.1.4. Application of the shifts

Application of the computed displacement vector (shifts) is trivial when the shifts are integer, since it corresponds to simple image translation and no interpolation is required. However when fractional shifts are applied there are multiple interpolation methods, based either on space interpolation (e.g., bilinear, bicubic) or on frequency domain interpolation (FFT-based). The choice of interpolation methods can lead to noticeably different results, a fact often overlooked. While frequency domain methods can be slower (since they require the computation of an inverse FFT), they tend to preserve more structure because they retain more frequency content of the signal and thus do not introduce any smoothing effects. For example, a rigid translation corresponds to a simple phase modulation in the frequency domain, which leaves invariant the power spectrum density of the image. Therefore, frequency interpolation also preserves the original SNR, as opposed to spatial interpolation methods that smooth the signal and increase the SNR. We discuss this issue in more detail in Section 3, where we show that frequency domain interpolation leads to crisper image statistics compared to spatial interpolation. Since spatial smoothing can also be achieved post-registration by default we use frequency domain interpolation. To preserve the dynamic range of the original data, the registered frame is restricted to take values between the minimum and maximum values of the original frame.

### 2.2. Evaluation metrics

Typically motion correction algorithms for calcium imaging data are evaluated on artificial datasets where known shifts are applied to registered data. On real data, evaluation typically occurs by visual inspection, where users observe the data (or a temporally downsampled version of it) before and after registration to assess the outcome of the registration. This makes the comparison of different algorithms on real datasets very hard and biased. In this paper we propose a series of simple metrics that can be used to quantify the performance of different algorithms. In section 3 we show that such metrics can be important for identifying locations where pw-rigid motion correction improves significantly upon simple rigid registration, a task very strenuous to be performed manually.

#### 2.2.1. Correlation with the mean metric

To evaluate the results of the motion correction algorithm across the different frames, we use a metric that is based on the similarity (pixel-wise, Pearson’s correlation coefficient *r*) between a reference template and each frame. For instance, one can compute for both the raw and corrected movie the correlation coefficient between each frame and the mean image across time, and then compare them. Intuitively, an increase in the correlation coefficient for a given frame indicates a better alignment with the mean^1^. To account for boundary effects during registration, a number of pixels around each boundary (e.g., equal to the maximum shift in each direction over time) is removed when computing the correlation coefficients.

This metric can be used to identify frames where the registration is successful or not, or to compare different motion correction algorithms at the level of individual frames. However, this metric critically depends on the smoothness properties of each frame which, as discussed in section3, can be affected by the method used to apply the computed displacements. In what follows, when using this metric we compare algorithms that register frames by applying shifts with the same method.

#### 2.2.2. Crispness and focus measures

An alternative measure is to quantify how crisp is a summary image before and after registration. This can be done by summing up the norm of the gradient field of the image on each location. If *I* is the resulting summary image then this measure of crispness can be defined as

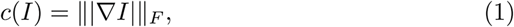

where *∇* denotes the gradient vector, *| · |* denotes the magnitude, and ∥ · ∥_*F*_ denotes the Frobenius norm. Examples of summary images include the mean image, or the correlation image (CI)^2^. Intuitively, a dataset with non-registered motion will have a blurred mean image, resulting in a lower value for the total gradient field norm. In addition to crispness, other measures of focus of the summarizing image can also be used. In this case as well, we expect that spatial interpolation methods can affect this measure, since introducing smoothing in each frame gives rise to a smoothed, higher valued correlation image with lower crispness.

#### 2.2.3. Residual motion quantification

To evaluate the performance of the algorithm, we can attempt quantifying motion before and after registration by using a different algorithm. In Section 3 we use the dense optical flow algorithm of Farnebäck [4] to estimate the residual motion and thus quantitatively evaluate the performance of the registration. In our setting, the algorithm estimates a motion field that attempts to match the current frame to the template. To do so, it relies on an efficient polynomial approximation of pixel neighborhoods to infer locally smooth displacement fields. In our hands, optical flow algorithms were particularly sensitive to the low/mid-SNR conditions of typical calcium imaging datasets. Therefore, in order to quantify the residual motion of other registration methods, the optical flow algorithm needed to operate on a downsampled version of the dataset to ensure robustness (and computational tractability). As such, we do not consider it as an appropriate method for registering calcium imaging data, but a useful and unbiased tool for assessing the performance of other methods.

### 2.3. Technical details

#### 2.3.1. Restricting maximum shifts

To avoid potential instabilities from corrupted or very sparsely labeled frames, the shifts allowed by the algorithm can be constrained within a user defined region. In practice, for each frame, NoRMCorre first computes the rigid displacement vector for the whole frame, with a user defined maximum allowed value, e.g. ∥**d**∥_∞_ ≤ *M*, where **d** is the rigid shift, *M* is the maximum allowed displacement in each direction, and ∥·∥_∞_ denotes the *l*_∞_ (max) norm. Then, the displacement vector for each patch is constrained within a given region centered around the rigid displacement vector, i.e., ∥**d**^*i*^ − **d**∥_∞_ ≤ *n*, where **d**^*i*^, is the displacement vector for patch *i*, and *n* is the maximum allowed deviation.

#### 2.3.2 Merging overlapping patches

To apply the shifts on overlapping patches we construct a set of weight interpolating functions that are used to ensure a smooth transition between registered neighboring patches. Consider the *i*-th patch, centered around the point (*x*_*i*_, *y*_*i*_) with size (*s*_*x*_, *s*_*y*_) and overlap (*o*_*x*_, *o*_*y*_), resulting in a total size (*s*_*x*_ + 2*o*_*x*_, *s*_*y*_ + 2*o*_*y*_). We define the trapezoid function

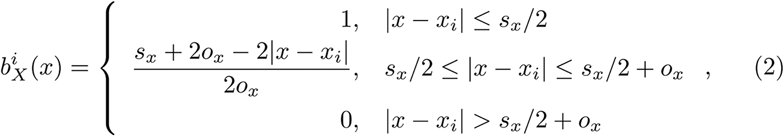

similarly the function 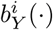 and the 2d function

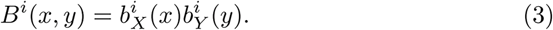

Then if *I*^1^*,…,I*^*K*^ are the reconstructed patches, extended to take values in the whole FOV, the interpolated registered frame is given by

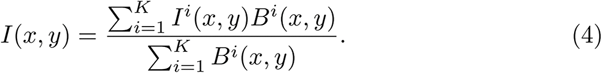

#### 2.3.3. Avoiding smearing by upsampling

When shifts among neighboring patches differ significantly, the interpolation explained above can introduce smearing effects. Take the case of two patches overlapping along the *x*-direction, whose *x*-shifts differ by exactly 1 pixel. When interpolating, the registered overlapping region will be simply a weighed average of two consecutive non-matching pixels along the *x*-direction, leading to a smeared result. Upsampling to a finer grid can alleviate this undesirable outcome. For example, if we up-sample the grid by a factor of 2, the difference in the displacements will be 0.5 pixels, thus inducing less smearing. We empirically observed that smearing occurs when shifts in overlapping patches differ by more than 0.5 pixels in either direction, and we suggest further upsampling to prevent it.

In theory, the grid could be upsampled to the point where each pixel has its own displacement vector. However, this approach can be computationally very slow, therefore introducing a trade-off between computational efficiency and smearing reduction. Hence, the upsampling factor can be chosen so that it fulfills the no-smearing condition with the following formula. If *n* denotes the maximum deviation from the rigid displacement for each patch, then two neighboring patches can have displacements that differ at most 2*n* pixels in each direction (an extreme case that in not expected to be encountered often in practice), and an upsampling factor of 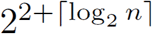, where ⌈*x*⌉ denotes the minimum integer greater or equal to *x*, guarantees the no smearing condition. For computational reasons, in practice we often use a smaller factor, and the interpolation is avoided for the frames where the smearing condition is not satisfied.

#### 2.3.4. Choosing patch size and amount of overlap

Our algorithm requires a template with strong reference points that facilitates robust matching and alignment. When splitting into patches to perform pw-rigid motion correction, each patch (together with its overlap) needs to contain enough signal to produce a clear template. In dark areas, for instance, it is difficult to find bright reference points and the alignment consequently fails. Empirically, for a typical 512 × 512 FOV with somatic imaging, an initial patch size of 128 × 128 (with additional 32 pixels of overlap in each direction) is a good choice. However, if the labelling is sparse then either a larger patch size and/or overlap might be required to ensure there is enough information for robust template alignment.

The amount of overlap between subpatches ensures a smooth interpolation between neighboring patches and alleviates boundary effects during the FFT registration. If *l* is the size of the initial patch along one dimension, *u* is the upsampling factor, and *M* + *n* is the allowed maximum displacement (maximum rigid displacement plus deviation), then we choose the overlap after upsampling to be larger than *M* + *n* − *l/u*, to ensure that each patch is not shifted by an amount larger than its dimension.

#### Software

Matlab code (also applicable to 3D volumetric imaging data) is available as a standalone package https://github.com/simonsfoundation/NoRMCorre. This package complements and will be integrated with the CNMF Matlab package for demixing and deconvolution of registered movies [13] available at https://github.com/epnev/ca_source_extraction. NoRMCorre is also implemented in Python https://github.com/simonsfoundation/CaImAn as part of the CaImAn package [5].

## 3. Results

We tested the algorithm on data collected *in vivo* with a two-photon microscope on a mouse expressing GCaMP6f in the parietal cortex, courtesy of S.A. Koay and D. Tank (Princeton University). The FOV had size 512 × 512 pixels and the data was acquired at 30Hz. Fig. 2 provides a demonstration of the performance of the rigid and piecewise rigid versions of our algorithm with respect to the various proposed metrics on a 2000 frame segment of the dataset. According to all the considered metrics, pw-rigid motion correction led to improved registration compared to plain rigid motion correction, which in turn improved significantly over the non-registered data. Fig. 2A shows a 100 × 100 pixel patch of the resulting mean for raw, rigid and pw-rigid corrected. By inspection, the pw-rigid correction preserves more fine structure, something that is also captured by the crispness metric (see eq. (1)) producing values of 4.3 × 10^3^, 6.69 × 10^3^, and 7.35 × 10^3^ for raw, rigid and piecewiserigid respectively. The same trend is also observed for the correlation with the mean metric (*Fig. 2b*) and the average per frame optical flow metrics (*Fig. 2c*), where the scatter plots demonstrate that the pw-rigid correction improves over the plain rigid correction for nearly all 2000 frames. Consistently, the optical flow metric shows that the improvement is also global in space (every region of the FOV exhibits less movement), with most of the remaining movement estimated to be around the boundaries and due to poorer SNR or other possible border effects (*Fig. 2e*). Fig. 2d shows the displacements along the *x*-axis for a small segment of frames (black), plotted against the displacements for each of the different patches (before upsampling). Connecting with Fig. 2b,c left, we notice that pw-rigid motion correction brings the most additional benefits over rigid motion correction when the dispersion of the displacements over the different patches is high, i.e., NoRMCorre estimates and corrects for a higher amount of non-rigid motion. The results are better displayed in movie format. Supplemental Movie 1 demonstrates the large variety of motion field patterns the algorithm estimates during the registration process. Supplemental Movie 2 shows a downsampled version of the results of rigid and pw-rigid registration, alongside the original data.

**Figure 2:**
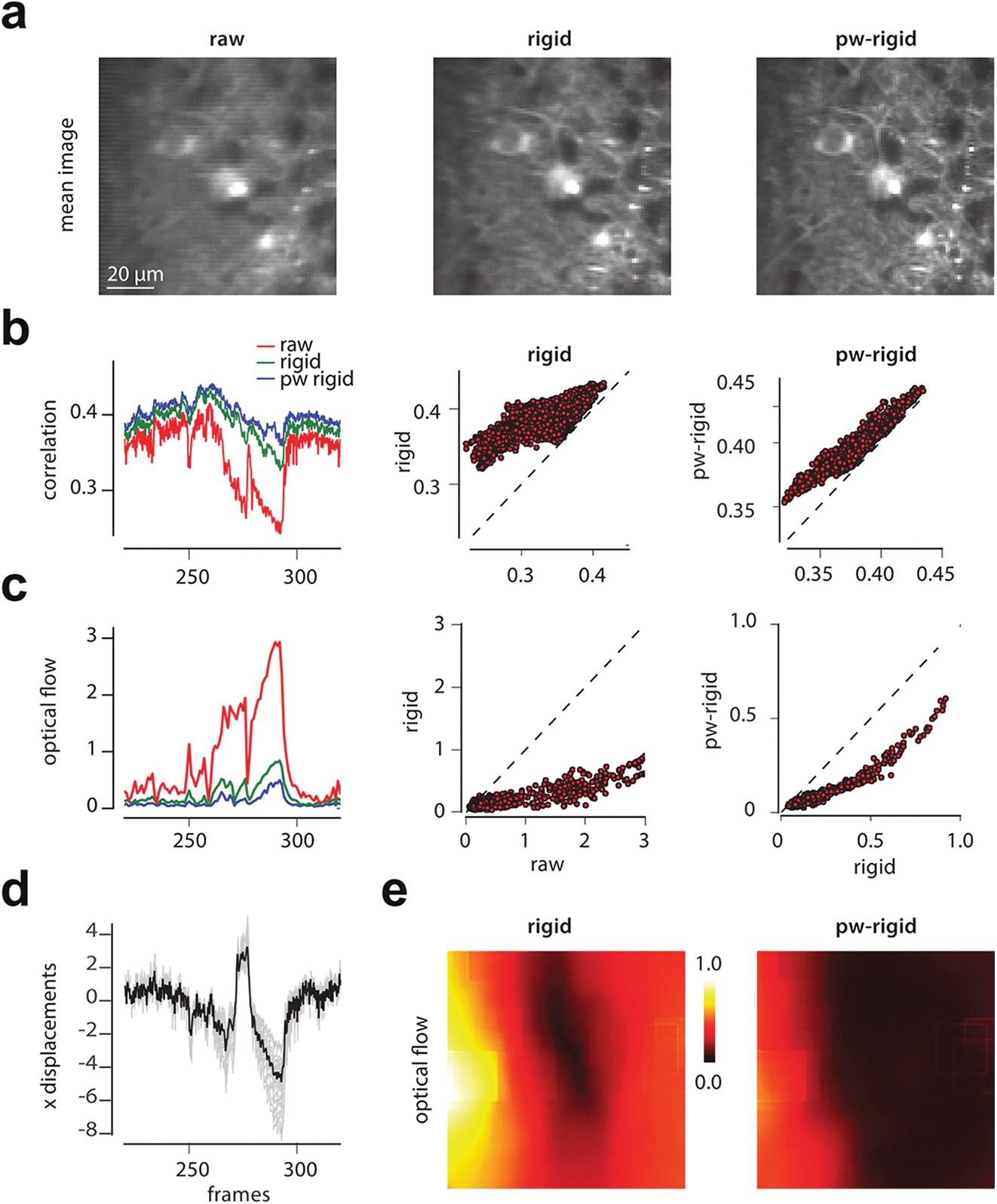
Illustration of performance on in vivo mouse parietal cortex data. *a*: Mean image of raw data (focused on a 100 × 100 pixels part of the FOV for clarity). Raw data (*left*), rigid corrected (*middle*) and piecewise-rigid corrected (*right*). NoRMCorre with pw-rigid correction results in a more structured mean image as quantified by the crispness of the image (1) (*c*(raw) = 4.3 × 10^3^*, c*(rigid) = 6.69 × 10^3^*, c*(piecewise) = 7.53 × 10^3^, measurements in absolute units). *b*: Quantification of performance based on the correlation with mean metric. For nearly every frame rigid correction improves over the raw data, and pw-rigid improves over rigid. *Left.* Mean correlation metric for a subset of frames. Scatter plot of frame-by-frame mean correlation metric of raw vs rigid (*center*) and rigid vs pw-rigid (*right*). *c, e*: Quantification of performance using the optical flow measure averaged over space (*c*, mean over space RMS value in pixels) and over time (*e*, mean over time RMS value of residual motion in pixels). Consistently, pw-rigid correction improves over plain rigid correction (*left*, frame by frame; *center*, scatter raw vs rigid; *right*, scatter rigid vs pw-rigid) and most of the remaining motion, as estimated with optical flow, remains on the boundaries of the FOV (*e, left*). *d*: Comparison of the rigid displacement (black) along the x-axis with the displacement of each patch for a subset of frames. The main benefits from the piecewise rigid correction, as can be seen from b and c (left) come at frames where the displacements exhibit maximum dispersion.

Next we compared NoRMCorre in its Python implementation with i) a Hidden Markov Model based algorithm [2], as implemented in the Python package SIMA [8], ii) the block-rigid approach of the Matlab package Suite2p [11], and iii) the Lucas-Kanade approach of Greenberg & Kerr [6]. These three methods are also suitable for non-rigid motion correction and have available implementations in Python (SIMA) or Matlab (Suite2p, Lucas-Kanade). We compared the three methods with respect to the quality metrics and the speed. For reference we also include the metrics of the non-registered data as well as the performance of rigid motion correction from the Python implementation of NoRMCorre. The results (*Table 1*) indicate that NoRMCorre achieves the best performance for crispness metrics and residual motion at a speed comparable to rigid motion correction, which is unsurprisingly the fastest method but produces the worst results in terms of residual motion. The residual motion was calculated with the dense optical flow (OF) algorithm of Farnebäck [4] in its OpenCV (v3.2, http://opencv.org) implementation, after temporal downsampling of the data to increase the SNR (see Section 2.2.3). We note that for all of the other three different methods, the best and reported results were obtained by taking blocks along the *x*-direction which is parallel to the raster scanning direction, demonstrating the fact that the largest part of the motion may not be due to raster scanning effect. Details of the various implementations are given in the supplement.

**Table 1:** Comparison of NoRMCorre with other non-rigid motion correction algorithm on a 2000 frame, 512 × 512 pixel *in vivo* mouse cortex dataset.

| 2000 frames | Crisp<br>(mean, a.u.) | Crisp<br>(CI) | Optical Flow<br>(RMS, pixels) | Time<br>(sec) | Interp.<br>Interp. |
| --- | --- | --- | --- | --- | --- |
| Original | 4301 | 9.87 | $1.574 \pm 1.502$ | — | — |
| Rigid | 6690 | 10.98 | $0.443 \pm 0.328$ | **40** | FFT |
| SIMA | 6678 | 9.25 | $0.244 \pm 0.082$ | 530 | Integer |
| Suite2p | 6694 | 9 | $0.246 \pm 0.08$ | 86 | FFT |
| Lucas-Kanade | 6394 | 10.36 | $0.197 \pm 0.084$ | 1856 | Bilinear |
| NoRMCorre | 7483 | 10.69 | $0.154 \pm 0.09$ | 89 | Bicubic |
| NoRMCorre | **7531** | **11.48** | **$0.15 \pm 0.09$** | 117 | FFT |

Table 1 also illustrates the effect of the interpolation method. When applying NoRMCorre with bicubic interpolation it achieves similar residual motion compared to NoRMCorre with Fourier interpolation albeit at a faster speed. However, the crispness of the mean and correlation images decreases due to the smoothness introduced by the bicubic interpolation. This point is highlighted even further in Fig. 3, where the correlation and mean images are shown for NoRMCorre and the Lucas-Kanade method, emphasizing the effect of different interpolation methods. Bilinear and bicubic interpolation smooths the data (*Fig. 3A, left and middle*), and biases upwards the correlation between neighboring pixels, as opposed to Fourier interpolation that retains the structure displayed by the weak correlations between neighboring pixels (*Fig. 3A, right*). On the other hand, the effect on the correlation with the mean metric is opposite leading to higher values for bilinear interpolation with Lucas-Kanade registration (0.499 ± 0.033), and bicubic interpolation with NoRMCorre registration (0.443 ± 0.017), as opposed to Fourier based interpolation with NoRMCorre which achieves a significantly lower value (0.399 ± 0.014). This highlights the sensitivity of this metric on the interpolation method, and why it should be used carefully in comparisons.

**Figure 3:**
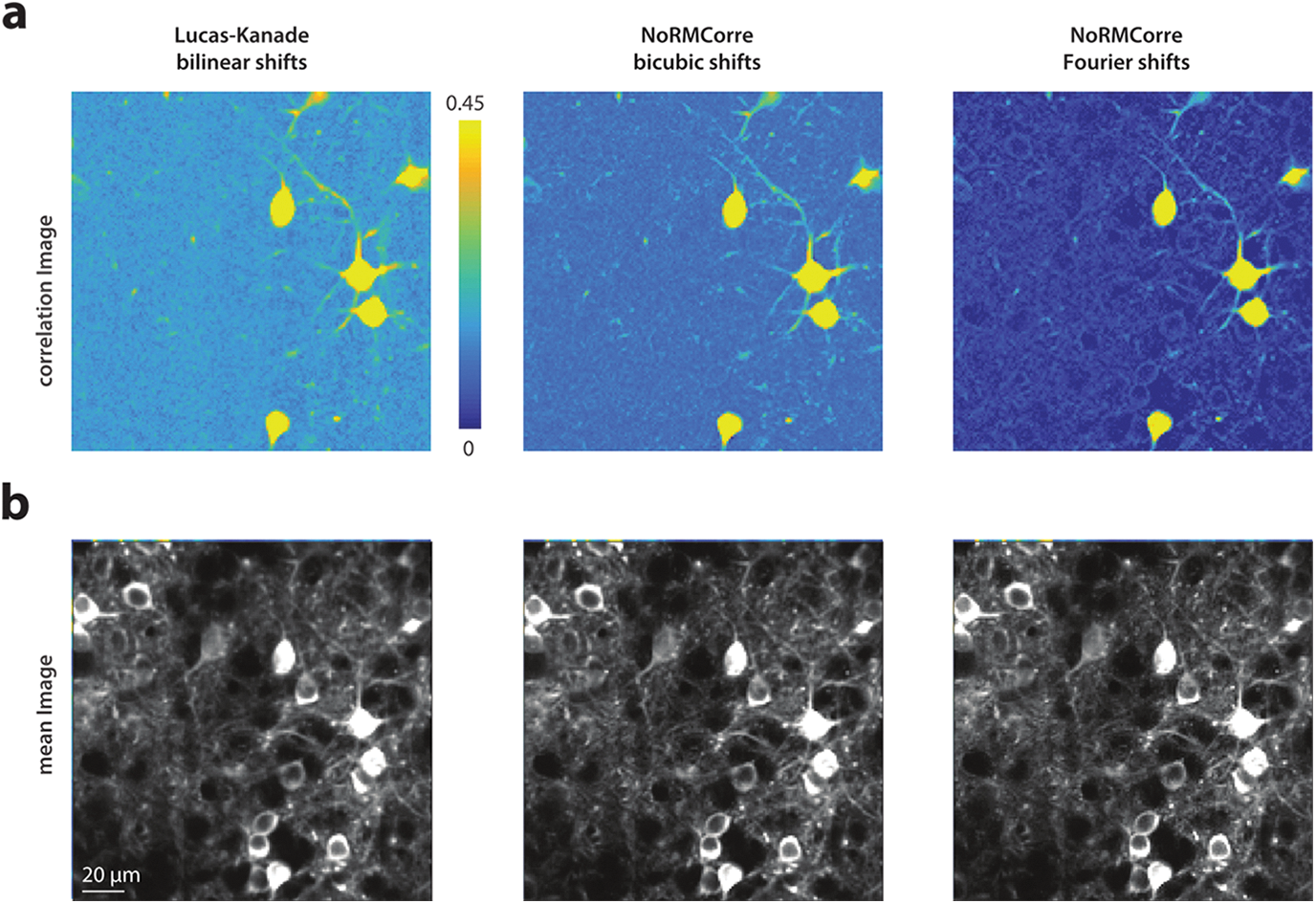
Effect of interpolation method on registered data. *a:* Correlation images for registered methods with 3 different methods restricted on a 170 × 170 pixels part of the FOV. Lucas-Kanade method with bilinear interpolation (*left*), and NoRMCorre with bicubic interpolation (*middle*) and Fourier interpolation (*right*). Fourier interpolation retains the weak correlation structure between neighboring pixels, whereas spatial interpolation “washes” away this structure by introducing smoothing during the shift application resulting in higher values for the correlation image. *b:* Mean images for the three approaches. The differences are less visible by eye, but quantitatively NoRMCorre with Fourier interpolation produced the crispest mean image (see *Table 1*).

## 4. Discussion

Non-rigid motion within a frame can occur not only due to slow raster scanning but also because of relative brain elastic deformation within the field of view. While faster raster scanning can result in higher imaging rates for a given FOV and thus reduce the amount of intra frame motion, modern methods enable imaging of even larger areas and/or volumes (e.g., Sofroniew et al. [14], Stirman et al. [15]) within which significant intra-frame motion is still possible. We believe that fast non-rigid motion correction will remain an important challenge in the future. NoRMCorre provides a simple and online method based on piecewise rigid template alignment that achieves state of the art results at a speed comparable to real time.

To better quantify the benefits of piecewise registration over rigid registration as well as to compare NoRMCorre with other non-rigid motion registration algorithms we developed some intuitive metrics that measure the crispness of the registered images and also used independent algorithms to estimate the amount of residual motion after registration. These metrics also highlighted the importance of the interpolation method that is chosen to apply the computed displacement vectors. While the effect of the smoothing introduced by the spatial interpolation methods might be minimal, and actually create the perception of a higher SNR, we took the stand that the statistics of the registered data should reflect the original input as much as possible, spatial smoothing can occur downstream in the analysis when necessary. We argued that by using the computationally more expensive Fourier based interpolation and avoiding any smoothing, one can better preserve the statistics of originally acquired data.

The ultimate goal of motion registration is to stabilize the FOV. This is important for segmentation reasons because several current source extraction methods identify sources by searching for groups of pixels that behave similarly with each other across time [13]. An alternative to such approaches would be to track individual neurons over time, an approach that has been taken when imaging freely moving *C. elegans* [10], where the deformations can be very dramatic. However, these methods tend to be computationally very expensive and have not yet found applications in registering other types of data.

The dataset used as an example in this paper pertains to two-photon, two-dimensional, raster scanning imaging of mostly cell bodies. However, our approach can also be applied to other types of imaging datasets. For the case of one-photon, microendoscopic data, high pass spatial filtering can be used to remove the bulk of the smooth background signal created by the large integration volume, and create stark reference points, prior to applying registration. NoRMCorre can also be readily applied to dense volumetric data (e.g., SCAPE microscopy [1]), where non-rigid motion can exist in all 3 directions. More details about such applications will be presented in the future.

## Acknowledgments

We thank D. Chklovskii, J. Magland (Flatiron Institute, Simons Foundation), J. Gauthier, S.A. Koay (Princeton University) and N. Sofroniew (Janelia Research Campus) for useful discussions. We thank S.A. Koay and D. Tank (Princeton University) for providing us with the *in vivo* mouse cortex dataset.

## Footnotes

1 If a static colored channel exists, these coefficients can in principle reach values very close to one, but in practice are limited by measurement noise. For variable channels their value is also limited by the time varying courses of the underlying neuronal processes.

2 The image where the value for each pixel is the average of the correlation coefficients between the pixel and its neighbors.

## Supplemental Material

### Dataset description

Data was obtained from the parietal cortex of a transgenic GCaMP6f-expressing mouse during a behavioral task. Field of view was approximately 500x500 *µm*^2^ (512 by 512 pixels) in size and at depth 125*µm* below the dura surface. Horizontal scans of the laser were performed using a resonant galvanometer, resulting in a frame acquisition rate of 30Hz.

### Implementation details

All analaysis and simulations were performed on a Dell Precision Tower 7910, 24 cores Intel(R) Xeon(R) E5-2643 v3 @3.40 GhZ, 128 GB RAM). For the pwrigid algorithm, the FOV was initially split in patches of size 128 × 128 pixels with an additional 32 pixels of overlap on each side. Each patch was further upsampled by a factor of 4. The algorithm was run in its offline mode with template obtained from the rigid registration of the first 500 frames. For the other three methods (SIMA, Suite2p, Lucas-Kanade), best results were obtained by taking blocks of size 512 × 16 pixels (excluding overlap), tiled horizontally (in parallel and not vertical to the raster scanning direction). For computation of the various metrics 12 pixels were removed from each side along both directions to avoid the boundary effects due to the registration. The optical flow algorithm was applied to a 5× downsampled version of the registered data to increase SNR and robustness.

### Description of Supplemental Movies

*Supplemental Movie 1. Depiction of the online pw-rigid motion correction procedure:.* Each frame of the original data (*top left*) is registered against a template (*bottom right*) in a piecewise rigid manner by shifting small patches according to the computed and upsampled motion field (*bottom left*). The resulting registered frame is shown on top right. Observance of the motion field shows the diverse non-rigid motion patterns that the algorithm estimates along both direction. The template is updated online during the registration process every 50 time steps. The movie is reproduced at the original data acquisition rate of 30 Hz.

*Supplemental Movie 2. NoRMCorre corrects for non rigid motion along both directions..* Comparison between the original data (*left*), corrected with rigid registration (*middle*) and piecewise rigid registration with NoRMCorre (*right*). Original and registered datasets are first downsampled 5× in time and then reproduced at 3× the original rate to aid the visual perception of the registration results. NoRMCorre with pw-rigid registration performs significantly better compared to rigid registration.

